# Increased FAN1 expression by mRNA-LNP attenuates CAG repeat expansion in Huntington patients’ iPSC-derived astrocytes

**DOI:** 10.1101/2023.11.24.568451

**Authors:** Yung-Chih Cheng, Gosia Nocula-Lugowska, Julita A. Ramirez, Xiaoyu Fan, Fang Jin, Zhihua Jiang, Eric Bennett, Jin Li, David Hokanson, Sneha Grandhi, Michelle Chen, Congsheng Cheng, Guan-Yu Lin, Laura Lin, Chris Lepsy, Javier Chaparro-Riggers, Laird Bloom, David Morrissey, Morag Stewart, Marija Tadin-Strapps, Shian-Huey Chiang

**Affiliations:** Target Sciences, Emerging Science & Innovation, Worldwide Research, Development & Medical; Pfizer Inc., 1 Portland St., Cambridge, MA, 02139, USA; BioMedicine Design, Worldwide Research, Development & Medical; Pfizer Inc., 1 Portland St., Cambridge, 02139, MA, USA; Global Science & Technology, Worldwide Research, Development & Medical; Pfizer Inc., 401 N Middleton Rd., Pearl River, NY, 10965, USA; BioMedicine Design, Worldwide Research, Development & Medical; Pfizer Inc., 1 Burtt Road, Andover, 01810, MA, USA; BioMedicine Design, Worldwide Research, Development & Medical; Pfizer Inc., 10777 Science Center Drive, San Diego, 92121, CA, USA; RNA Accelerator, Emerging Science & Innovation, Worldwide Research, Development & Medical; Pfizer Inc., 1 Portland St., Cambridge, MA, 02139, USA; RNA Medicine, Centers for Therapeutic Innovation, Emerging Science & Innovation, Pfizer Inc., 450 East 29^th^ St., New York, NY, 10016, USA

## Abstract

Expansion of repeat sequences within the human genome can lead to disease pathogenesis, such as Huntington’s Disease, primarily affecting the nervous system. Genome-wide association studies (GWAS) of age-at-onset in Huntington’s disease (HD) patients demonstrated DNA mismatch repair (MMR) genes are modifiers of somatic expansion and may be potential therapeutic targets for repeat expansion (RE) disorders. FAN1, a Fanconi anemia-associated nuclease, has been reported as an influencer of repeat expansion in the RE mouse models. Here, we show the first demonstration that FAN1 knock-out in HD patient-derived fibroblasts and results in increased CAG repeat length. We also develop a robust novel cell-based platform using stem cell technology to produce the HD patients’ iPSC-derived astrocytes (iAstro). This platform is a disease-relevant system and has a significantly wider assay window, making it more suitable to assess the effect of gene modulation on CAG repeats. A substantial and exponential increase in repeat instability was exhibited in this HD patient’s iPSC-derived astrocytes platform. Over-expression of FAN1 protein via *FAN1* plasmid transfection in this platform reduced CAG repeat instability, suggesting that upregulation of FAN1 protein may have a potential protective effect in CAG repeat expansion for a therapeutic setting. We leveraged the mRNA-LNP modality to enhance FAN1 protein expression and revealed that codon-optimized *FAN1* mRNA-LNP robustly prevented increased CAG repeat in HD patients’ iPSC-derived astrocytes platform. The data from these cell-based platforms highlight that FAN1 plays a protective role in attenuating expanded somatic *HTT* CAG repeats and shed light on new therapeutic directions against repeat expansion disorders.

## INTRODUCTION

Repeat expansion (RE) disorders are due to unstable repetitive sequences in the human genome, either in the coding or non-coding regions, and usually affect the nervous system’s normal function and lead to severe diseases. Currently, there are over 40 human genetic disorders known to be caused by these expansions in DNA sequence. These are categorized as RE disorders, such as Huntington’s disease (HD), Fragile X syndrome (FXS), Myotonic Dystrophy (DM1), Spinocerebellar Ataxia (SCA), and Frontotemporal Dementia/Amyotrophic Lateral Sclerosis (FTD/C9-ALS, GC in C9orf72). The clinical manifestation of RE disorders appears earlier and becomes more severe with aging as the diseases are inherited from one generation to the next. Evidence shows a strong correlation between the size of the repeat expansion and the age at onset and/or the severity of the disorders (1). Huntington’s disease is a dominant, monogenic, and inherited neurological disorder, which is driven by the expansion of CAG repeat sequence (≥ 40 CAG repeats) within exon 1 of the huntingtin (*HTT*) gene, resulting in an expanded polyglutamine tract in the huntingtin protein (2,3). Mutant HTT protein disrupts transcription, interferes with mitochondrial function, and is further aberrantly modified post-translationally to involve complex pathogenic mechanisms (4). Expansion of CAG repeats occurs in distinct somatic and meiotic tissues, but the neurodegenerative pathology is primarily due to the massive loss of neurons in the striatum and cortex tissues (2). This selective neuronal loss occurs preferentially in medium-spiny neurons of the striatum, and this damage then massively extends to other brain regions (5). Besides neuronal cells, the majority of cells in the brain are glia, which support the survival and function of neurons. Astrocytes are one of the main types of glia and express glutamate transporters that uptake extracellular glutamate to prevent glutamate neurotoxicity (6). It has also been identified that mutant HTT is expressed in glial cells in the brains of HD mouse models and in HD patients (7,8), although how the mutant HTT generated by CAG expanded repeats in glia drives neuropathology and massive neuronal loss is still unclear.

Emerging evidence from genome-wide associated study (GWAS) from HD patients suggests the age-at-onset of HD is highly correlated with genes in the DNA mismatch repair (MMR) pathway, including MLH1, FAN1, PMS1, MSH3, PMS2, and LIG1 (9). Several studies also demonstrated in mouse models that deficiency of MSH3, MLH3, PMS2, and MLH1 presents a protective effect to prevent somatic repeat expansion in the repeat expansion mouse models (10-13). However, knocking out FAN1 (FANCD2 and FANCI-associated nuclease 1) exacerbated acceleration of somatic CAG repeat expansion in the HD mouse model and CGG repeat expansion in the FXS mouse model (14-16) highlighting the opposite direction of FAN1 protein from other MMR proteins, in the repeat expansion regulation. Overexpression of FAN1 protein in the U2OS (osteosarcoma) cell line expressing mutant HTT exon 1 provides stabilization of expanded HTT CAG repeats (17) by binding MLH1 protein to restrict MLH1 recruitment by MSH3 or through its nuclease activity to execute an accurate repair (18). The data is consistent with analyses linking FAN1 loss-of-function variants with earlier onset of Huntington’s disease (19), suggesting that FAN1 nuclease protein can modify the repeat expansion rate and prevent error-prone repair from suppressing CAG repeat expansion.

Current therapeutic approaches for Huntington’s disease focus on targeting the mutant HTT production pathway, including HTT transcription and the translation of its mRNA, to alleviate pathogenesis. However, a previous study has shown the length of the longer CAG repeat tract is the primary determinant of the age at disease onset (20). It demonstrated that other genetic factors could modify disease occurrence and timing. Given the exacerbated effect from knocking out FAN1 in the HD mouse model and the stabilized effect from overexpression of FAN1 in the *in vitro* U2OS RE cellular model, the strategy for enhancing FAN1 protein expression could be a potential therapeutic approach to prevent somatic CAG repeat expansion and slow the disease progression. Following the success of the COVID-19 vaccine, mRNA-based therapeutics have become a potential new drug class in a broad range of therapeutic applications, including immunotherapeutic, protein-replacement, and regenerative medicine (21). One advantage of mRNA technology is the ability to increase the endogenous expression of a target protein without modifying the genome. In addition, lipid nanoparticle (LNP) delivery of RNA cargo has recently been shown to be a well-tolerated and safe for systemic delivery system of genetic payloads to target tissues (22,23). Thus, leveraging the mRNA-LNP platform to enhance FAN1 protein expression may represent a new therapeutic approach to stabilize somatic CAG repeat expansion and instability in Huntington’s disease.

Here, we have employed two cell-based models to study the role of FAN1 protein in regulating CAG repeat length. Besides HD patient-derived fibroblasts, we have also developed a novel Repeat Expansion (RE) *in vitro* cellular model by leveraging the HD patient iPSC platform. This system provides an advantage over the other currently available *in vitro* RE cellular models as it is CNS-relevant and more translatable to a disease setting. We have leveraged both models to study the effect of modulation of FAN1 protein levels on CAG repeat expansion. Knockout of FAN1 in HD patient-derived fibroblasts exacerbates CAG repeat expansion, while over-expression FAN1 stabilizes CAG repeat length in HD patient iPSC-derived astrocytes. To test the modulated effect of FAN protein *in vivo*, we formulated *FAN1* mRNA-LNP to assess FAN1’s influence on CAG repeat expansion in the liver, a peripheral tissue that presents the highest CAG variability in post-mortem and HD mice samples (24,25). Systemic administration of a single dose of codon-optimized *FAN1* mRNA-LNP demonstrated a 130-fold increased level over normal FAN1 protein expression at 6 hr, though the expression was not sustained beyond 24 hr. It suggests that additional mRNA-LNP optimization is needed to achieve therapeutically relevant levels of FAN1 protein *in vivo*. Briefly, this finding demonstrates the function of FAN1 protein in preventing the repeat expansion within our established HD patient iPSC-derived astrocytes, providing a therapeutic direction for RE disorders.

## MATERIALS AND METHODS

### Cell lines, reagents, and services

Primary Huntington (HD) patient fibroblasts (GM09197; CAG = ∼ 180/18) were purchased from Coriell Institute. Fibroblasts cells were maintained in EMEM (Eagle’s Minimum Essential Medium) (Sigma-Aldrich; M4655) supplemented with 15% non-heat inactivated FBS (Fetal bovine serum) (Sigma-Aldrich; F0926), MEM Non-Essential Amino Acid (Gibco; 11140-050), and 100 IU Penicillin/100 μg/mL Streptomycin (Corning; 30-001-Cl). Huh7 (Human hepatocellular carcinoma) cells were grown in DMEM (Dulbecco’s Modified Eagle Medium) (Gibco; 11885-084; Low glucose), and HepG2 (Human hepatocellular carcinoma) cells were grown in EMEM (ATCC; 30-2003). All culture mediums were supplemented with 10% heat-inactivated FBS (Gibco; 10082-147), MEM Non-Essential Amino Acid (Gibco; 11140-050), and 100 IU Penicillin/100 μg/mL Streptomycin (Corning; 30-001-Cl). Hepa 1-6 (Murine hepatoma) cells were grown in DMEM (Gibco; 11995-065) supplemented with 10% heat-inactivated FBS (Gibco; 10082-147), 0.15% Sodium Bicarbonate (Gibco; 25080-094), and 100 IU Penicillin/100 μg/mL Streptomycin (Corning; 30-001-Cl). SB431542 (TGF-β/SMAD inhibitor; 1614) and ROCK inhibitor (1254) was from Tocris. LDN193189 (BMP inhibitor; 04-0074-02) was from Reprocell. Heparin sodium salt, Dextrose, DMSO (Dimethyl sulfoxide), and other chemicals were from Sigma-Aldrich unless otherwise specified. TaqMan probes were all from Life Technologies. Recombinant IL-1β and TNF-α proteins were from R&D Systems. Human IL-6 ELISA kit was from Life Technologies. FAN1KO HD patient fibroblasts were generated and received from Synthego. Mouse genotyping and HTT CAG number were analyzed by Laragen. HTT CAG repeat expansion and instability analysis was also performed by Laragen.

### HD patient iPSC-derived astrocytes differentiation and culture

HD patient iPSCs (ND50036) were obtained from NINDS and maintained in mTeSR Plus (STEMCELL Technologies; 100-0276) medium on Matrigel (Corning; 354277) coated plate. The differentiation protocol of HD patient iPSC-derived astrocytes is based on the published protocol from Stem Cell Reports (26). Briefly, dual SMAD inhibition (0.1 μM LDN193189 and 10 μM SB431542) was applied to differentiate HD patient iPSCs to Neural rosette formation in N2B27 medium (DMEM/F-12-GlutaMax (Gibco; 10565-042), 1% N2 (Gibco; 17502-048), 2% B27 minus vitamin A (Gibco; 12587-010)) supplemented with MEM Non-Essential Amino Acid (Gibco; 11140-050), and 100 IU Penicillin/100 μg/mL Streptomycin (Corning; 30-001-Cl) and then into NPCs (Neural Progenitors) in NPC medium (N2B27 medium with supplements and with additional 20 ng/mL FGF2 (Gibco; PHG0021) and 1 ug/mL Laminin (Gibco; 23017-015)). Split NPCs 1:3 every week with accutase (STEMCELL Technologies; 7920). NPCs were further differentiated into human astrocytes by seeding dissociated single cells at 15,000 cells/cm^2^ density on Matrigel-coated plates in Astrocyte medium (ScienCell; 1801). Cells were fed every 2 days and split to the initial seeding density (15,000 cells/cm^2^) with accutase every week for 30 – 90 days for astrocyte maturation.

### Optimization of mouse FAN1 coding sequence

For codon optimization, sequences were assigned a score based on a linear combination of underlying metrics. One metric was the codon adaptation index (CAI) (27), with codon normalization performed in Table S1. Another metric was “rolling” CAI, where the CAI is calculated over all possible windows of 4 consecutive codons, and the reported value for the metric is the lowest value observed for any single window. Another metric was the fraction of unpaired bases in the MFE secondary structure of the encoding mRNA, as calculated using default settings in ViennaRNA (28). The last metric considered was the fraction of bases in the MFE secondary structure that were involved in stretches of more than 30 consecutive paired bases (“Long run fraction”). The optimization used multiple stages employing either a Monte Carlo genetic algorithm (MC/GA) or steepest descent search (SD).

For Monte Carlo searches, a single parent sequence was chosen as the seed, and mutated child sequences were generated by randomly mutating codons to synonymous codons according to a specified probability (the mutation rate; Table S2 & S3). When making a mutation, synonymous replacement codons were chosen according to the weights in Supporting Table S1. The child sequences constituted a single generation of the genetic algorithm. The mutated child sequence with the best metric value was chosen to seed a successive generation. The mutation rate decreased by a certain amount (mutation decay rate) with each generation. After a certain number of generations (Table S2 &S3), the optimization stopped, and a set of winning sequences was chosen such that optimal metric scores were balanced against primary sequence diversity. More specifically, the chosen sequences had the best metrics that could be obtained while ensuring no two chosen sequences had more than a specified limit of primary sequence identity. These sequences were fed into the next cycle of optimization. Each cycle used a fixed number of processor cores, with the potential input sequences from the previous round divided evenly among the processors. Repeats of more than 6 of the same nucleotides in a row were disallowed, as were various restriction sites.

Steepest descent searches involved stepping through the sequence one codon at a time from N to C terminus, trying all synonymous codons, and choosing the codon giving the optimal metric value. The optimization metric was always minimized. Because it is hypothesized that CAI should be maximized to increase expression, CAI and rolling CAI were generally given negative weights. Two variations on the above optimization scheme were used. The optimization schedule for Method 1 is given in Supporting Table S2. In the first round, a reverse-optimization is performed, to generate a sequence with poor CAI (poor predicted expression). This process removes any bias in the input sequence and ensures that random searching can productively explore many regions of sequence space without being stuck in a local minimum. Method 1 is designed to give a diverse panel of sequences for testing. The optimization schedule for Method 2 is given in Supporting Table S3. While similar to Method 1, Method 2 is designed to give the single sequence with highest CAI. It does not consider rolling CAI or secondary structure. Because it would tend to choose the highest frequency codon for each amino acid, repeated amino acid motifs would lead to repeated nucleic acid motifs, which might be problematic. Therefore, an additional filter was applied to Method 2: the sequence was not allowed to contain any repeated stretches of 10 bases anywhere in the sequence. Both methods were applied to mouse FAN1 (NCBI Reference Sequence NM.177893.4) to generate optimized candidates for testing.

### Generation of human/mouse *FAN1* plasmids and mouse *Fan1* mRNA-LNP

The DNA encoding human *FAN1* (NCBI Reference Sequence: NM.014967.5 and NP.055782.3) mouse *Fan1* (NCBI Reference Sequences: NP.808561 and NM.177893.4) or a codon optimized mouse *Fan1* was either used for the direct transfection of cells or was used as a template for RNA synthesis. Codon optimized sequences were designed using the CAI method described above (Codon Adaptation Index) (27). Standard mammalian expression vector was used for direct cell transfection. The DNA template used for *in vitro* mRNA synthesis had additional elements such as 5’ and 3’ untranslated sequences (UTRs) and a poly-A tail. Messenger RNA was synthesized with uridine substituted with N1-methylpseudouridine. Mature RNA product had an N7-methylated cap to improve translation efficiency (29). RNA-LNP formulations were prepared using microfluidic mixing with Precision Ignite NanoAssemblr (Precision Nanosystems, Vancouver, BC). In brief, lipids were dissolved in ethanol at molar ratios of 46.3:42.7:9.4:1.6 (ALC-0315:cholesterol:DSPC:ALC-0159) as previously described (30,31). Lipid mixture was combined with a 50 mM citrate buffer (pH 4.0) containing mRNA at a ratio of 3:1 (aqueous: ethanol). LNPs were dialyzed overnight at 4°C into 10 mM Tris (pH=7.5) buffer using a 10k MWCO (molecular weight cut off) dialysis cassette. Resulting formulations were concentrated using Amicon Ultra Centrifugal Filter devices with a 30 kDa cut-off (EMD Millipore, Billerica, MA) and sterile filtered using 0.22 μm PES filters (Fisher Scientific, NY, USA). 300 mM sucrose was added to the final formulation. All LNPs were tested for particle size, RNA encapsulation, RNA integrity and endotoxin (< 1 EU/ml of endotoxin).

### Recombinant mouse FAN1 protein expression and purification

Cell pellet (80 g) from 4 L of HEK293 cells transfected with muFAN1_His8_HA expressing vector was resuspended in 400 mL of cold Lysis buffer (TBS, pH 7.5, 20 mM Imidazole, 1 mM TCEP, 1 mM MgCl_2_, 3 μL Benzonase, EDTA-free C0mplete tablets, 0.05% Triton X-100) and lysed by Microfluidizer, lysate was centrifugated at 25,000 x g for 45 minutes to pellet cellular debris and insoluble material. Supernatant was decanted and batch bound with Ni-Excel Sepharose resin and eluted with 250 mM Imidazole resulting in 10 mg of protein. muFAN1_His8_HA was applied to a Superdex200 16/60 SEC column pre-equilibrated and run in SEC buffer (PBS, 1 mM TCEP, 1 mM EDTA) resulting in 2 mg of final protein.

### FAN1 plasmid and mRNA-LNP transfection in HD patient iPSC-derived astrocytes

To over-express FAN1 protein in HD patient iPSC-derived astrocytes, we leveraged human *FAN1* (*hFAN1*) plasmid and *mFan1* mRNA-LNP by transfection. HD patient iPSC-derived astrocytes were transiently transfected with 2 μg *hFAN1* plasmid and control plasmid (pTT5 vector) by Lipofectamine 3000 (Invitrogen; L3000008) every week. Similarly, 15 μg/mL *mFan1* mRNA-LNP was transfected directly into HD patient iPSC-derived astrocytes every week. Cell pellets were collected every week for CAG repeat instability analysis.

### Mouse breeding and maintenance

All procedures performed on animals were in accordance with USDA regulations, conducted in compliance with the *Guide for the Care and Use of Laboratory Animals* (National Research Council), and reviewed and approved by Pfizer’s Institutional Animal Care and Use Committee. *Htt*^*Q111*^ HD Knock-in mice (*Hdh*^*Q111*^) were maintained on a *C57BL/6J* background by breeding with *C57BL/6J* mice from The Jackson Laboratory (Bar Harbor, ME). Genotyping of *Hdh*^*Q111*^ mice was performed with genomic DNA extracted from tail or ear biopsies at weaning or in adult tissue samples obtained at necropsy and was carried out by Laragen. All animals were housed under specific pathogen free and controlled temperature conditions, in individually ventilated caging with free choice food and water and 12:12 hour light/dark cycles.

### Treatment of *Hdh*^*Q111*s^mice with *mFan1* mRNA-LNP

Native *mFan1* mRNA-LNP and codon-optimized *mFan1* mRNA-LNP were prepared by Pfizer internally. 8-week-old *Hdh*^*Q111*^ mice (CAG 105 - 115 males) received 2 mg/kg codon-optimized *mFan1* modRNA-LNP, 2 mg/kg native *mFan1* modRNA-LNP, or LNP formulation buffer by single tail vein injection. Three (native *Fan1* control) or 4 animals (CO#5.2, LNP buffer control) per group were euthanized and tissues harvested at 6 hours post-injection, and the remaining animals in each group were euthanized with tissue harvest at 24 hours post-injection. Two sections of liver were snap frozen in liquid nitrogen for evaluation of FAN1 protein or mRNA levels, respectively.

### Statistical analysis

Statistical analyses were performed using GraphPad Prism v.9. Data are presented as mean values ± standard deviation using One-Way ANOVA, with Bonferroni comparison test. The instability index of CAG repeats in *Hdh*^*Q111*^ allele in mouse tissues or *HTT* allele in HD patient fibroblasts or iPSC-derived astrocytes was quantified as a published study (32).

## RESULTS

### Knocking out *FAN1* in HD patient-derived fibroblasts increases HTT somatic CAG repeat expansion

One of the current therapeutic approaches for Huntington’s disease aims to delay age at onset or disease progression of this devastating disease (33-35). A previous GWAS study revealed multiple genetic modifiers at gene loci suggesting DNA integrity maintenance may be a potential strategy to modify the age-at-onset of Huntington’s disease. Among those genes, FAN1, at the chromosome 15 locus, was identified as one of the modifier genes that influences HD age at onset (9,19); suggesting that FAN1 plays a crucial role in regulating repeat expansion.

To test the functional significance of FAN1 on repeat expansion, we knocked out *FAN1* in HD patient fibroblasts (GM09197; CAG = ∼ 180/18) by gene editing. There was approximately 50% KO efficiency in *FAN1* KO pool of HD patient fibroblasts. After three passages of culture (3 weeks; passage every week), compared with MOCK control, *FAN1*KO HD patient fibroblasts strikingly increased the level of accumulated CAG expansion index (Fig. 1a). The CAG size and the repeat instability index were also increased in HD patient fibroblasts after knocking out *FAN1* (Fig. 1b and 1c), with no noticeable difference in the cell viability. It is the first demonstration of the effect of *FAN1* knockout in HD patient derived fibroblasts.

**Figure 1.**
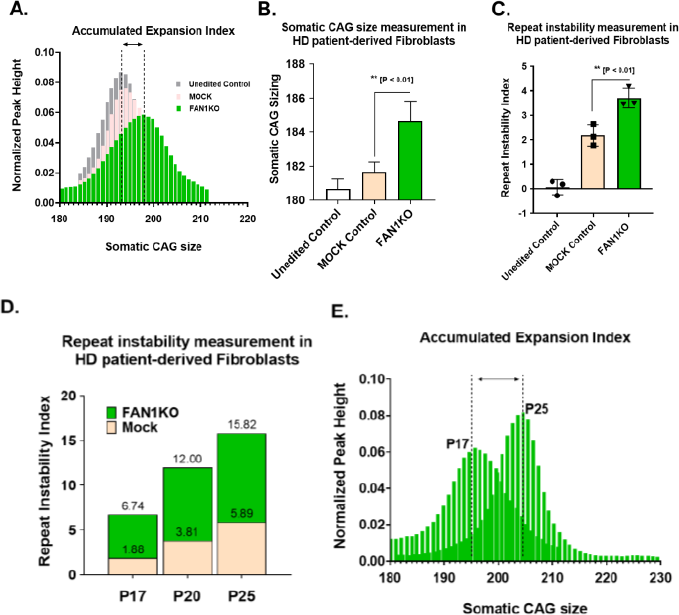
Loss of *FAN1* increases *HTT* CAG repeat instability in HD patient-derived fibroblasts. **A)** Representative plot of *HTT* CAG size distribution in HD patient-derived fibroblasts (CAG 180). **B)** *HTT* CAG repeat size in control and *FAN1*KO HD patient-derived fibroblasts. **C)** Quantification of CAG repeat instability in HD patient-derived fibroblasts. **D)** Potently increased *HTT* CAG repeat instability in *FAN1*KO HD patient-derived fibroblasts following different passaging times. **E)** Augmented expansion of *HTT* CAG size in HD patient-derived fibroblasts after 5 times of passaging.

As a surrogate for the aging process, we wanted to understand whether the CAG repeat in HD patient-derived fibroblasts expands during the time in culture and thus we cultured MOCK and *FAN1*KO HD patient fibroblasts from Passage 17 (P17) to 25 (P25). The CAG repeat instability index increased in MOCK HD patient fibroblasts from 1.88 to 5.89 after eight weeks of culture. Significantly, the CAG repeat instability index rose to 15.82 in *FAN1*KO HD patient fibroblasts (Fig. 1d). The level of accumulated CAG expansion index and CAG size markedly increased in *FAN1*KO HD patient fibroblasts after long-term culturing (Fig. 1e). These observations strongly imply that FAN1 plays a role as a negative regulator in CAG repeat in this HD patient-derived cell type. The findings are consistent with increased somatic instability of CAG repeat due to longer mutant HTT *in vivo* (24,36).

### Generation of HD astrocytes from patient-derived iPSC

While HD patient-derived fibroblasts are a feasible cell-based model that can interrogate the role of the DNA mismatch repair (MMR) gene on CAG repeat expansion, the assay window is often rather narrow for a screening mode. Moreover, connecting the disease phenotype with the modulated effect of MMR protein from this fibroblast platform is challenging. We thus sought to develop a cell-based model that may be more disease-relevant and potentially offer a wider assay window to study the effects of MMR gene modulation, even viable for screen mode. Glial cells are primary neuron-supporting cells in the nervous system. Astrocytes, one of the main glial cells, closely contact neurons, blood vessels, and other glial cells to support neuronal function. Astrocytes are implicated to be highly involved in central nervous system (CNS) disorders (37-39). Astrocytes from HD mouse model display a significant variety in the molecular and cellular aspects relative to control (38,40). Previous reports also offer multiple differentiation protocols to derive human astrocytes from human iPSC or ESC, in an attempt to recapitulate human primary astrocytes (26,41-43) for studying CNS disorders. To set up an *in vitro* nerve-cell model relevant to HD, we applied an efficient protocol from Julia *TCM et al*. (26) to differentiate human iPSC into functional astrocytes. We generated neural progenitors from HD patient-derived iPSC (NDS00143; ND50036; CAG = ∼ 109). Utilizing this differentiation protocol consisted of three stages. The first two-week-long stage drives human iPSC into neural rosettes to be regionally patterned and then differentiated into neural progenitors (NPCs) (Fig. 2a). The second stage is the gliogenic switch, which transition NPCs from neurogenic to gliogenic lineage. While remaining proliferative, these cells express features consistent with an immature astrocytic phenotype (44,45) (Fig. 2a) which is responsive to signals driving terminal differentiation and maturation. The last stage is the maturation of human astrocytes. Immature proliferative astrocytes can be induced to spontaneously differentiate by astrocyte medium (ScienCell) (Fig. 2a) with low initial seeding density (nearly single cells: 15,000 cells/cm2) and requires protracted culture durations for astrocyte maturation (> 90 days). Mature astrocytes were confirmed with an average composition of 95% S100β+ and 90% GFAP+ cells by immunocytochemistry and quantified by Operetta High Content Analysis (HCA) system (Fig.2b). Compared with the origin hiPSC line, human OCT4 (POU5F1; hiPSC marker) expression was not expressed in the induced human astrocytes (Fig. 2c). However, as with primary astrocytes, there was substantial variability in mature astrocyte markers. S100β was compatible with the expression in primary astrocytes (Fig. 2d), but GFAP expression was robustly lower in the induced human astrocytes, although demonstrated to be highly induced after glial progenitors (26) (Fig. 2e).

**Figure 2.**
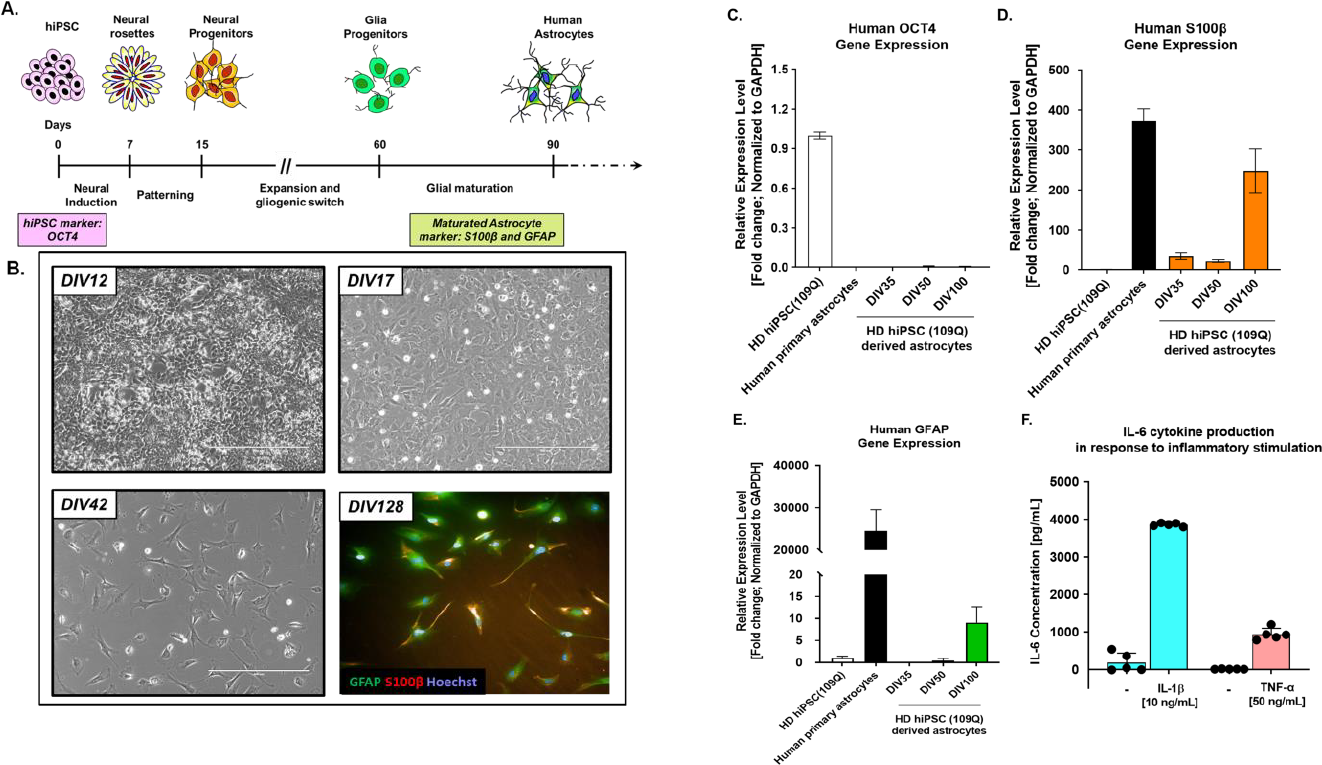
Modeling and characterization of HD patient iPSC-derived astrocytes. **A)** HD patient iPSCs are initially converted into neural rosettes and then regional patterned to neural progenitors. Neural progenitors can be expanded in the presence of astrocyte growth factors. Longterm expansion of neural progenitors allows the gliogenic switch toward matured astrocytes. **B)** Morphological changes during the differentiation process. Maturated astrocytes are expressing astrocyte-specific marker GFAP (green) and S100β (red). Cell-type specific marker expression of HD iPSC and matured astrocytes, measured by qRT-PCR. **C)** hiPSC associated marker: OCT4. Human primary astrocytes were used as a positive control. Increasing **D)** S100β and **E)** GFAP over the maturation of astrocytes. **F)** Pro-inflammatory cytokine production in response to inflammatory stimulation in human iPSC-derived astrocytes. Successfully differentiated human astrocytes respond to stimulation by IL-1β (10 ng/mL) and TNF-α (50 ng/mL) by secreting IL-6.

Astrocytes are immunocompetent cells that regulate neuroimmune response and actively participate in neuroinflammation by secreting cytokines and chemokines that are protective or detrimental to neuronal function, survival, and diseases (46,47). To confirm generation of fully functional, immuno-competent astrocytes, the inflammatory cytokine, IL-6, was measured in the induced human astrocytes after stimulation with IL-1β (10 ng/mL) or TNF-α (50 ng/mL) for 24 hrs. The total IL-6 in the non-stimulated condition was low (Fig. 2f), indicating the induced human astrocytes were in the non-reactive status. However, we observed a high level of IL-6 production and high cell survival rates following IL-1β (10 ng/mL) or TNF-α (50 ng/mL) treatment. This result suggests that like primary human astrocytes, human iPSC-derived astrocytes are reactive to inflammatory stimuli and immunocompetent in further sustaining inflammation by producing the pro-inflammatory cytokine (IL-6).

### CAG repeats are unstable in HD patient iPSC-derived astrocytes

Huntington’s disease (HD) is caused by a CAG repeat expansion in the somatic tissues in HD patient brains (9,48), and these repeats could be unstable in dividing cells (glial cells) and non-dividing cells (neurons) (49,50). To examine whether there is an unstable repeat expansion/instability in our developed *in vitro* RE cellular model, HD patient iPSC-derived astrocytes, cell lysates were collected at different time points during differentiation for the repeat instability calculation. We observed a significant increase of CAG size and a robust expanding from accumulated expansion index in HD patient iPSC-derived astrocytes compared to HD patient iPSC. Intriguingly, the increased CAG size continued to expand in this HD astrocytes from DIV (Day of *In Vitro*) 72 to DIV 130 (Fig. 3a). Besides, there was a tremendous exponentially increase of repeat instability in HD patient iPSC-derived astrocytes after repeat instability calculation (Fig. 3b). Our result showed that unstable repeats occur in HD patient iPSC-derived astrocytes, providing an advantageous platform for studying repeat expansion disorders, especially in Huntington disease.

**Figure 3.**
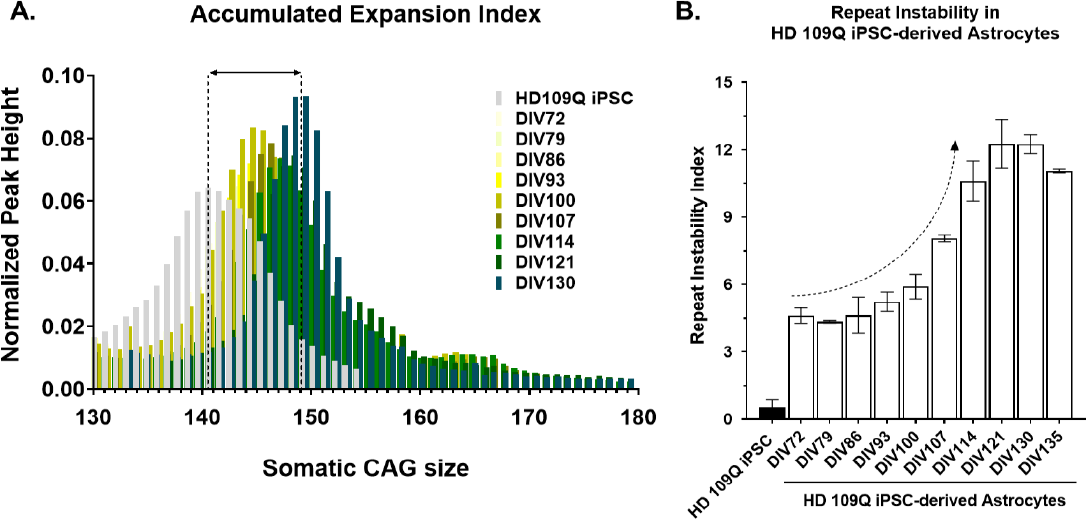
*HTT* CAG expansion in HD patient hiPSC-derived astrocytes. Fragment analysis showed the change in modal *HTT* CAG repeat size over time in HD patient iPSC-derived astrocytes. b) Quantification of *HTT* CAG repeat instability in HD patient iPSC-derived astrocytes revealed dramatically increase in *HTT* CAG instability in culture.

### FAN1 overexpression delays CAG repeat expansion in HD patient iPSC-derived astrocytes

Recent evidence demonstrates increasing FAN1 expression presented a stabilized effect in the *HTT* CAG repeat in U2OS *HTT* exon 1 cell line to slow CAG expansion rate (17). To verify FAN1 protein’s impact in the regulation of repeat expansion, we used a human *FAN1* plasmid to overexpress FAN1 protein in our *in vitro* RE platform built from HD patient iPSC-derived astrocytes. After multiple transfections (once per week), the repeat instability in HD iPSC-derived astrocytes did not raise as quickly as the control, instead showing a slow elevation (Fig. 4a). We also observed the *HTT* CAG size, and the accumulated expansion index were not increased as much as the pTT5 and iAstro only control (Fig. 4b). Although there may be an influence from multiple transfections on decreased repeat instability in pTT5 control group, overexpression of FAN1 protein still demonstrated a robust protective effect to stabilize *HTT* CAG repeat instability and to prevent CAG repeat expansion in HD iPSC-derived astrocytes. We also harvested cell lysates after several transfection cycles to further examine FAN1 protein expression. FAN1 protein was significantly increased in the HD iPSC-derived astrocytes at DIV107 after three cycles of transfection, with the protein expression levels 15-fold higher than the control group (Fig. 4c). The result indicated overexpression of FAN1 protein efficiently prevents CAG repeat instability and repeat expansion in this HD iPSC-derived astrocytes cellular model.

**Figure 4.**
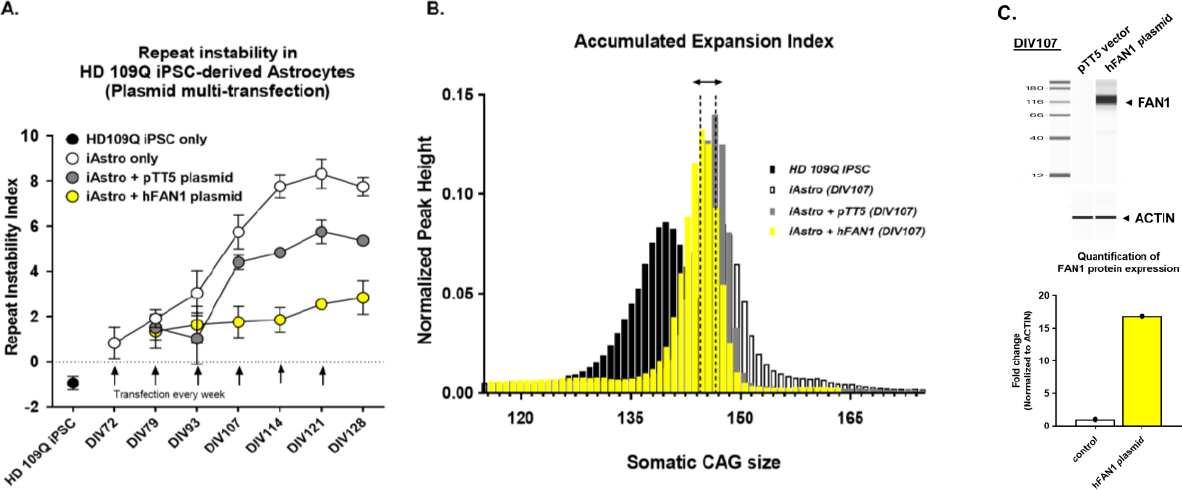
Overexpression FAN1 protein expression prevents *HTT* CAG repeat expansion. **A)** Quantification of *HTT* CAG repeat instability in HD patient iPSC-derived astrocytes. Overexpression FAN1 protein expression prevented HTT CAG repeat instability increase in HD patient iPSC-derived astrocytes after multiple times of transfection with human *FAN1* plasmid. **B)** Fragment analysis traces *HTT* CAT repeat size distribution over 8 weeks. Ectopic expression of human FAN1 protein reduces *HTT* CAG repeat size in HD patient iPSC-derived astrocytes. **C)** Cell lysates harvested from HD patient-derived astrocytes at DIV107 (day *in vitro*). Immunoblots developed with FAN1 and ACTIN antibodies are shown. Quantification of FAN1 protein expression at DIV107. Overexpression of FAN1 protein by multiple transfections of human *FAN1* plasmid increased over 10 folds of protein expression in HD patient-derived astrocytes.

### Production and Optimization of FAN1 mRNA-LNP to enhance FAN1 protein expression

HD murine models have been widely used to study either *HTT* somatic CAG repeat expansion or pathogenesis of disease (51,52). *Hdh*^*Q111*^ mouse model, which has 111 CAG repeat inserted in the first exon of mouse Htt gene, demonstrates the highest CAG repeat instability in liver and striatum tissues (32), allowing us to study the effect of FAN1 in the regulation of repeat expansion *in vivo*. To augment FAN1 protein expression in the HdhQ111 mouse model, we also intended to leverage mRNA-LNP platform, which provides an easy means to overexpress target protein in animal models, particularly the liver tissue (53). We first synthesized the wild-type (WT) coding sequence of murine *Fan1* (native; NV) and generated several codon-optimized sequences of murine *Fan1* with a goal of enhancing FAN1 protein expression. Native (NV) *mFan1* mRNA sequence, six codon-optimized *mFan1* mRNA sequences, and one mutant *mFan1* mRNA sequence that removed two putative degradation signals (D-KEN box to prevent ubiquitination pathway (54)) were tested in an expi293 cell line. The initial testing in expi293 cell line confirmed four codon-optimized sequences (#1, #2, #4, #5.2) increased protein expression 3-4 fold, relative to native sequence, at 24 hrs after mRNA transfection. The increased FAN1 protein expression lasted for 72 hrs after mRNA transfection. Significantly, the protein expression level in codon-optimized *mFan1* (#1 and #5.2) was 8.6-fold higher than native *mFan1* at 72 hrs after mRNA transfection (Fig. S1). We then continued testing these four codon-optimized *mFan1* DNA constructs in HepG2 cell line to validate their protein expression in human liver cell line. The four codon-optimized *mFan1* DNA constructs (#1, #2, #4, and #5.2) all increased FAN1 protein expression at 48 hrs after transfection compared with control (Opti-MEM only) and native *mFan1* coding sequence (Fig. 5a). Notably, #5.2 codon-optimized *mFan1* sequence showed a significant increase in protein expression starting at 24 hrs after transfection. Furthermore, we generated #4 and #5.2 codon-optimized *mFan1* mRNAs and examined their efficiency of translation in FAN1 protein expression in Huh7 (human) and Hepa1-6 (murine) liver cell lines after mRNA transfection. As expected, #5.2 codon-optimized *mFan1* mRNA presented clear effectiveness in showing a robust increase of FAN1 protein expression at 24 hrs after mRNA transfection in Huh7 and Hepa1-6 cell lines (Fig. 5b). Quantification data also indicated there was an ∼ 15-fold induction of FAN1 protein expression in Huh7 cell line after transfection with #5.2 codon-optimized *mFan1* mRNA. Likewise, a ∼ 13-fold induction of FAN1 protein expression after transfection with #4 codon-optimized *mFan1* mRNA (Fig. 5c) was observed. Based on the *in vitro* validation, #5.2 codon-optimized *mFan1* mRNA was selected as the premier candidate for subsequent FAN1 protein overexpression studies.

**Figure 5.**
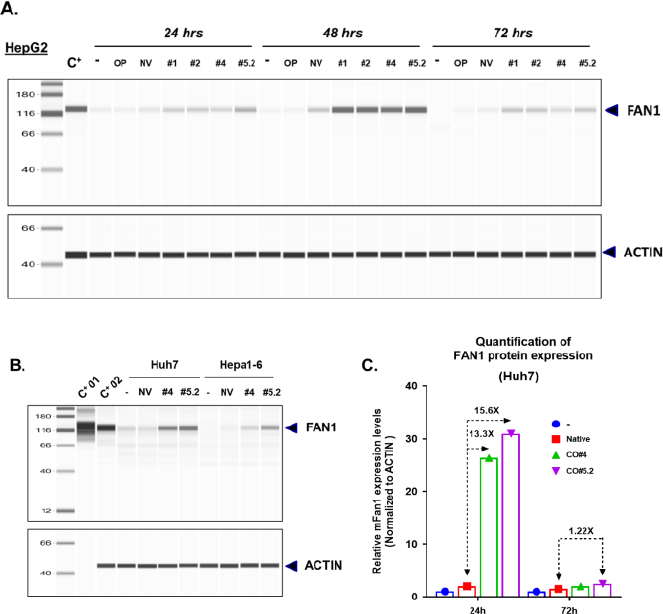
*In vitro* characterization and validation of WT and codonoptimized mouse *Fan1* mRNAs in mammalian liver cancer cell lines. **A)** Capillary electrophoresis immunoblotting for detecting FAN1 protein expression in HepG2 human liver cancer cell line transfected with control (Opti-MEM only), plasmid constructs of native (NV) coding sequence or different codon-optimized coding sequences *(#1, #2, #4, and #5*.*2)* encoding murine FAN1 proteins. Positive control (C^+^): recombinant mouse FAN1 protein. **B)** 24 hrs after transfection with mRNA of native (NV) coding sequence or codon-optimized coding sequences (*#4 and #5*.*2*) encoding murine FAN1 protein in Huh7 (human) and Hepa1-6 (mouse) liver cancer cell lines. **C)** Quantification of FAN1 protein expression in Huh7 human cancer cell line after transfection with codon-optimized mouse *Fan1* mRNAs.

### Efficacy of codon-optimized *mFan1*_#5.2_ mRNA-LNP in preventing increased CAG repeat instability in HD iPSC-derived astrocytes

To assess the efficacy of codon-optimized *mFan1*_#5.2_ mRNA in FAN1 protein re-expression and in regulating repeat expansion, an *in vitro* validation was performed by conducting multiple transfections of codon-optimized *mFan1*_#5.2_ mRNA-LNP in HD iPSC-derived astrocytes. The transfections were performed twice weekly as duration of FAN1 expression from mRNA-LNP delivery was shown to be 72 hrs (Fig. 5), to boost higher FAN1 protein expression, relative to endogenous FAN1 protein expression. There was a marked reduction in repeat instability in HD iPSC-derived astrocytes after multi-transfection with codon-optimized mFan1#5.2 mRNA-LNP (Fig. 6a). Importantly, continuous overexpression of FAN1 protein not only slowed the increased repeat instability but also resulted to some contraction of CAG repeat in HD iPSC derived-astrocytes after multiple-transfections with codon-optimized *mFan1*_#5.2_ mRNA-LNP (Fig. 6a & 6b). The results are consistent with earlier reports and prove FAN1 is critical in regulating CAG repeat expansion, although the detailed molecular machinery is still unclear (17,18,55,56).

**Figure 6.**
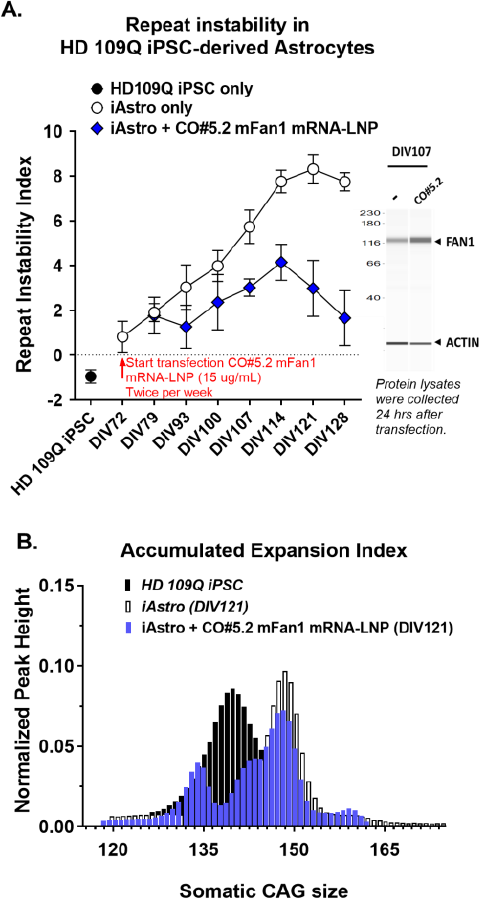
Multiple transfections with codon-optimized *mFan1* mRNA-LNP increases FAN1 protein expression and reduces HTT CAG repeat expansion in HD patient-derived astrocytes. **A)** Quantification of *HTT* CAG repeat instability in HD patient iPSC-derived astrocytes. Overexpression FAN1 protein by multiple transfections with codon-optimized *mFan1* mRNA-LNP (CO#5.2) prevented HTT CAG repeat instability increase in HD patient iPSC-derived astrocytes. **B)** Fragment analysis traces *HTT* CAT repeat size distribution over 8 weeks. Ectopic expression of FAN1 protein reduces *HTT* CAG repeat size in HD patient iPSC-derived astrocytes.

### Codon-optimized *mFan1*_#5.2_ mRNA-LNP administration transiently boosts FAN1 protein expression *in vivo*

Given the effective activity of codon-optimized *mFan1*_#5.2_ mRNA-LNP in HD iPSC-derived astrocytes, we aimed to further explore enhanced *in vivo* expression of FAN1 protein expression and efficacy in prevention of repeat expansion in the HD mouse model. The first dosing regimen tested the durability of codon-optimized *mFan1*_#5.2_ mRNA-LNP. Codon-optimized *mFan1*_#5.2_ mRNA-LNP was dosed in an 8-week-old male HD mouse model (*Hdh*^*Q111*^) and liver tissue harvested for analysis of protein expression at different time points (Fig. 7a). Interestingly, enhancement of FAN1 protein expression using codon-optimized *mFan1*_#5.2_ mRNA-LNP was substantial up to 6 hrs after administration, but not at 24 hrs (Fig. 7b). Compared with native mFan1 mRNA-LNP or LNP formulation buffer control, codon-optimized *mFan1*_#5.2_ mRNA-LNP generated over 100-fold higher FAN1 protein at 6 hrs post-dose; however, FAN1 protein returned to base line by 24 hrs after dosing (Fig. 7c). This short duration of protein expression was somewhat surprising as we have previously showed longer duration of expression (57) (ABCB4 and unpublished data) using similar mRNA-LNP platform. The short duration of FAN1 protein expression will require frequent repeat administrations to reduce CAG expansion in HD murine models, thus increasing the challenge for *in vivo* evaluation. Further studies are needed to understand the mechanisms underlying the short half-life of FAN1 protein and perform necessary platform optimization to enable lower dosing frequency.

**Figure 7.**
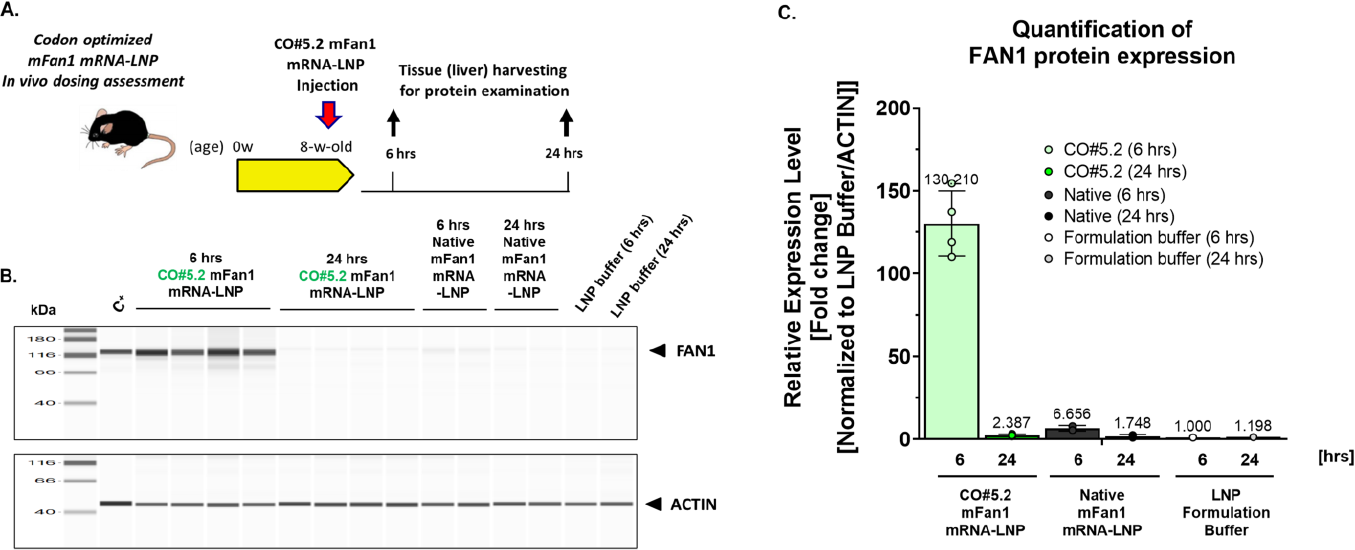
*In vivo* validation of codon-optimized *mFan1* mRNA-LNP in FAN1 protein expression. **A)** Scheme of experimental design of *in vivo* validation of codon-optimized *mFan1* mRNA-LNP (CO#5.2). 8-week-old male *Hdh*^*Q111*^ mice received single dosing of CO#5.2 *mFan1* mRNA-LNP (2 mg/kg, i.v.), native *mFan1* mRNA-LNP, or LNP formulation buffer (n=4 for each time point). **B)** Capillary electrophoresis immunoblotting for detecting FAN1 protein expression in liver tissue harvested at 6 hrs and 24 hrs after dosing. Positive control (C^+^): recombinant mouse FAN1 protein. **C)** Quantification of FAN1 protein expression in mouse liver tissues at different time points after dosing with codon-optimized mouse *Fan1* mRNA-LNP, native *mFan1* mRNA-LNP, and LNP formulation buffer.

## DISCUSSION

Current therapeutic approaches of Huntington’s disease include preventing CAG repeat expansion to delay disease progression. RE mouse models, e.g., HD mouse or FXN mouse models, have been exploited in previous studies to identify and validate potential modifiers to regulate repeat expansion. However, these mouse models take months to conclude since RE is an age-dependent pathology phenotype. Consequently, having the ability to quickly validate potential gene modifiers that can regulate the kinetics of repeat expansion progression associated with RE disorders in a cell-based assay would be very valuable. Here, we describe developing a new RE cell-based system by leveraging stem cell technology to differentiate HD patient iPSC into astrocytes, a cell type in the CNS is highly correlated to the pathology of RE disorders (58-60). Our HD iPSC-derived astrocyte platform displays the expanded CAG repeats within six weeks and is characterized by a robust/exponential increase of repeat instability (Fig. 3) compared to HD patient-derived fibroblasts and other RE cell-based assays. Although historical studies suggest a strictly neuronal pathology in patient’s brains due to triplet repeat expansions, other groups have demonstrated that the expression of Htt with expanded CAG repeats in mouse astrocytes manifest functional atrophy (52). In addition, a recent study indicates white matter glial cells in CNS of myotonic dystrophy type 1 (DM1) patients present unstable and longer repeats (61) relative to healthy. Thus, the overall weight of evidence shows astrocytes are involved in repeat expansion disorders. Our RE cell-based platform using HD patients’ iPSC-derived astrocytes provides a practical system for screening potential genetic modifiers for the therapeutics of RE disorders.

FAN1, a Fanconi anemia-associated nuclease, was identified as a crucial element for DNA interstrand crosslink repair in the Fanconi anemia (FA) pathway (62-64). Previous researches indicate FAN1 is involved in the DNA mismatch repair (MMR) pathway (65) and displays onset-hastening or delaying haplotypes of HD (19). In this study, we knocked out FAN1 in HD patient-derived fibroblasts, showing the exaggerated effect in CAG repeat instability. Then, we demonstrated that overexpression of FAN1 protein in this HD patient iPSC-derived astrocytes model for the first time, using DNA plasmid transfection, provided a protective effect in prevention of increased CAG repeat expansion and instability in these HD iPSC-derived astrocytes, a CNS relevant *in vitro* cell-based platform. Our result demonstrated that FAN1 is a negative regulator of triplet repeat expansion. Overexpression of FAN1 protein in HD patient iPSC-derived astrocytes effectively slowed CAG repeat instability and presented a robustly protective effect. These findings are consistent with the previous studies (17,18) in U2OS osteosarcoma cell line expressing mutant *HTT* exon 1, although the detailed mechanism by which exogenous FAN1 overexpression regulated repeat instability is still unclear.

We also leveraged mRNA therapy focusing on FAN1 to regulate repeat expansion in HD patient iPSC-derived astrocytes. The basic principle of mRNA therapeutics is to deliver mRNA into the target cell, where cellular machinery translates the mRNA into a functional protein. Thus, mRNAs can be widely applied across vaccine development, protein replacement therapy, and the treatment of genetic disorders with an efficient and safe delivery approach. Although we have successfully demonstrated the functional delivery of our *FAN1* mRNA-LNP into human/mouse liver cell lines and our HD patient-derived astrocytes, the durability of FAN1 protein expression *in vivo* remains to be optimized. Notwithstanding elevated expression of FAN1 for up to 6 hours following *in vivo* transfection, FAN1 protein levels declined rapidly, as evidenced by a swift decay of FAN1 protein in liver tissues after administration, despite using the same codon-optimized *FAN1* mRNA-LNP *in vitro* (Fig. 5-7). FAN1 protein has been proved as a mitotic substrate of the anaphase-promoting cyclosome complex (APC/C), an E3 ubiquitin ligase complex to degrade cell cycle-related proteins during G1 phase. Since FAN1 is regulated by APC/C and is degraded during the mitotic exit, the duration of FAN1 expression is essential during mitotic progression (54), though it’s activity thereafter remains unclear. This degradation pathway may explain why the expression of FAN1 protein by mRNA-LNP is only elevated in mouse liver tissue out to 6 hrs, but not at 24 hrs, after dosing *in vivo*. Further studies are needed to verify this hypothesis. In addition, it may be worth evaluating FAN1 expression using new generation of mRNA platforms, e.g., saRNA (self-amplifying RNA) or circRNA (circular RNA), which may extend protein expression, enable more durable expression *in vivo*, and provide clinically feasible and relevant dose regimens. This result reveals the different biological features of FAN1 in different cell types and exposes the imperfection of our *in vitro* HD cell-based platform, set apart from live organisms.

In summary, we leveraged Huntington patients’ iPSC-derived astrocytes to establish a reliable repeat expansion cell-based assay to examine the function of FAN1 protein in preventing repeat expansion/instability using a novel mRNA-based therapeutic system. Our results suggest FAN1 overexpression may have protective effects in a therapeutic setting. The new generation of mRNA therapy may provide a new therapeutic direction for repeat expansion disorders.

## ACKNOWLEDGMENTS

This study was supported and sponsored by Pfizer. We thank the BioMedicine Design (BMD) team (Aaron D’Antona, Amy Tam, Chong Wang, Divya Patel, Dokyong Kim, Edward Lavallie, Jessica Min-DeBartolo, Joshua Lee, Justin Cohen, Martina Zafferani, Mikaela Palandra, Ruiting Lin, Wayne Stochaj) for the preparation, generation, and QC of the hFAN1 and mFAN1 constructs and mRNA-LNP, Joanne Brodfuehrer, Hendrik Neubert, Nick Psychogios and Roberto Ortiz for pharmacokinetics analyses, Pragya Rampuria for fruitful discussions, and Annette Sievers for AAV QC support; Global Science & Technology (GST) and Comparative Medicine (CM) team (Michael Cinque, Tariel Turner, Ryan Misonznick, Reyna Prosnitz, Carlos Hall) for their efforts with the *in vivo* studies; Dr. Uwe Schoenbeck, Dr. Morten Sogaard, Dr. David Shields, Dr. Brian Bates, Dr. Paul Wes, Dr. Benedikt Bosbach, Dr. Brooke Coni Trousdale and other Pfizer colleagues in Emerging Science & Innovation (ES&I) for their support and stimulating discussion in this project. We are also grateful to Dr. Christine Bulawa, Dr. James Fleming, Dr. Claudia Huichalaf, Dr. Amrutha Pattamatta, Dr. Jian An in Rare Disease Research Unit (RDRU) for the experimental discussion and insights on the repeat expansion programs.

## AUTHOR CONTRIBUTION

Y.-C.C., G.N.-L., J.A.R., X.F, M.S., S.-H.C., M.T.-S. contributed to the conception and design of the study. Y.-C.C., G.N.-L., J.A.R., F.J., Z.J., J.L, D.H., M.C., C.C., G.-Y.L. performed the experiments and analyzed the data. E.B. performed the codon-optimized mRNA design. S.G. performed the genetic study. L.L., C.L., J.C.-R., L.B., D.M. provide oversight of this program. Y.-C.C., G.N.-L., J.A.R., X.F, M.S., S.-H.C., M.T.-S. interpreted the results of the experiments. Y.-C.C. prepared the figures and drafted the manuscript. Y.-C.C., G.N.-L., D.M., J.A.R., M.S., M.T.-S. edited and revised the manuscript. All authors reviewed the manuscript and approved submission of the final version. This research work was supported by Pfizer.

## REFERENCES

1. Andrew, S.E., Goldberg, Y.P., Kremer, B., Telenius, H., Theilmann, J., Adam, S., Starr, E., Squitieri, F., Lin, B., Kalchman, M.A. et al. (1993) The relationship between trinucleotide (CAG) repeat length and clinical features of Huntington’s disease. Nat Genet, 4, 398–403.

2. MacDonald, M.E., Barnes, G., Srinidhi, J., Duyao, M.P., Ambrose, C.M., Myers, R.H., Gray, J., Conneally, P.M., Young, A., Penney, J. et al. (1993) Gametic but not somatic instability of CAG repeat length in Huntington’s disease. J Med Genet, 30, 982–986.

3. (1993) A novel gene containing a trinucleotide repeat that is expanded and unstable on Huntington’s disease chromosomes. The Huntington’s Disease Collaborative Research Group. Cell, 72, 971–983.

4. Podvin, S., Rosenthal, S.B., Poon, W., Wei, E., Fisch, K.M. and Hook, V. (2022) Mutant Huntingtin Protein Interaction Map Implicates Dysregulation of Multiple Cellular Pathways in Neurodegeneration of Huntington’s Disease. J Huntingtons Dis, 11, 243–267.

5. Vonsattel, J.P. and DiFiglia, M. (1998) Huntington disease. J Neuropathol Exp Neurol, 57, 369–384.

6. Sood, A., Preeti, K., Fernandes, V., Khatri, D.K. and Singh, S.B. (2021) Glia: A major player in glutamate-GABA dysregulation-mediated neurodegeneration. J Neurosci Res, 99, 3148–3189.

7. Jing, L., Cheng, S., Pan, Y., Liu, Q., Yang, W., Li, S. and Li, X.J. (2021) Accumulation of Endogenous Mutant Huntingtin in Astrocytes Exacerbates Neuropathology of Huntington Disease in Mice. Mol Neurobiol, 58, 5112–5126.

8. Jansen, A.H., van Hal, M., Op den Kelder, I.C., Meier, R.T., de Ruiter, A.A., Schut, M.H., Smith, D.L., Grit, C., Brouwer, N., Kamphuis, W. et al. (2017) Frequency of nuclear mutant huntingtin inclusion formation in neurons and glia is cell-type-specific. Glia, 65, 50–61.

9. Genetic Modifiers of Huntington’s Disease Consortium. Electronic address, g.h.m.h.e. and Genetic Modifiers of Huntington’s Disease, C. (2019) CAG Repeat Not Polyglutamine Length Determines Timing of Huntington’s Disease Onset. Cell, 178, 887–900 e814.

10. Dragileva, E., Hendricks, A., Teed, A., Gillis, T., Lopez, E.T., Friedberg, E.C., Kucherlapati, R., Edelmann, W., Lunetta, K.L., MacDonald, M.E. et al. (2009) Intergenerational and striatal CAG repeat instability in Huntington’s disease knock-in mice involve different DNA repair genes. Neurobiol Dis, 33, 37–47.

11. Pinto, R.M., Dragileva, E., Kirby, A., Lloret, A., Lopez, E., St Claire, J., Panigrahi, G.B., Hou, C., Holloway, K., Gillis, T. et al. (2013) Mismatch repair genes Mlh1 and Mlh3 modify CAG instability in Huntington’s disease mice: genome-wide and candidate approaches. PLoS Genet, 9, e1003930.

12. Wheeler, V.C., Lebel, L.A., Vrbanac, V., Teed, A., te Riele, H. and MacDonald, M.E. (2003) Mismatch repair gene Msh2 modifies the timing of early disease in Hdh(Q111) striatum. Hum Mol Genet, 12, 273–281.

13. Hayward, B.E., Steinbach, P.J. and Usdin, K. (2020) A point mutation in the nuclease domain of MLH3 eliminates repeat expansions in a mouse stem cell model of the Fragile X-related disorders. Nucleic Acids Res, 48, 7856–7863.

14. Zhao, X., Lu, H. and Usdin, K. (2021) FAN1’s protection against CGG repeat expansion requires its nuclease activity and is FANCD2-independent. Nucleic Acids Res, 49, 11643–11652.

15. Zhao, X.-N. and Usdin, K. (2018) FAN1 protects against repeat expansions in a Fragile X mouse model. DNA Repair, 69, 1–5.

16. Loupe, J.M., Pinto, R.M., Kim, K.H., Gillis, T., Mysore, J.S., Andrew, M.A., Kovalenko, M., Murtha, R., Seong, I., Gusella, J.F. et al. (2020) Promotion of somatic CAG repeat expansion by Fan1 knock-out in Huntington’s disease knock-in mice is blocked by Mlh1 knock-out. Hum Mol Genet, 29, 3044–3053.

17. Goold, R., Flower, M., Moss, D.H., Medway, C., Wood-Kaczmar, A., Andre, R., Farshim, P., Bates, G.P., Holmans, P., Jones, L. et al. (2019) FAN1 modifies Huntington’s disease progression by stabilizing the expanded HTT CAG repeat. Hum Mol Genet, 28, 650–661.

18. Goold, R., Hamilton, J., Menneteau, T., Flower, M., Bunting, E.L., Aldous, S.G., Porro, A., Vicente, J.R., Allen, N.D., Wilkinson, H. et al. (2021) FAN1 controls mismatch repair complex assembly via MLH1 retention to stabilize CAG repeat expansion in Huntington’s disease. Cell Rep, 36, 109649.

19. Kim, K.H., Hong, E.P., Shin, J.W., Chao, M.J., Loupe, J., Gillis, T., Mysore, J.S., Holmans, P., Jones, L., Orth, M. et al. (2020) Genetic and Functional Analyses Point to FAN1 as the Source of Multiple Huntington Disease Modifier Effects. Am J Hum Genet, 107, 96–110.

20. Lee, J.M., Ramos, E.M., Lee, J.H., Gillis, T., Mysore, J.S., Hayden, M.R., Warby, S.C., Morrison, P., Nance, M., Ross, C.A. et al. (2012) CAG repeat expansion in Huntington disease determines age at onset in a fully dominant fashion. Neurology, 78, 690–695.

21. Sahin, U., Kariko, K. and Tureci, O. (2014) mRNA-based therapeutics--developing a new class of drugs. Nat Rev Drug Discov, 13, 759–780.

22. Polack, F.P., Thomas, S.J., Kitchin, N., Absalon, J., Gurtman, A., Lockhart, S., Perez, J.L., Perez Marc, G., Moreira, E.D., Zerbini, C. et al. (2020) Safety and Efficacy of the BNT162b2 mRNA Covid-19 Vaccine. N Engl J Med, 383, 2603–2615.

23. Adams, D., Gonzalez-Duarte, A., O’Riordan, W.D., Yang, C.C., Ueda, M., Kristen, A.V., Tournev, I., Schmidt, H.H., Coelho, T., Berk, J.L. et al. (2018) Patisiran, an RNAi Therapeutic, for Hereditary Transthyretin Amyloidosis. N Engl J Med, 379, 11–21.

24. Wheeler, V.C., Auerbach, W., White, J.K., Srinidhi, J., Auerbach, A., Ryan, A., Duyao, M.P., Vrbanac, V., Weaver, M., Gusella, J.F. et al. (1999) Length-dependent gametic CAG repeat instability in the Huntington’s disease knock-in mouse. Hum Mol Genet, 8, 115–122.

25. Telenius, H., Kremer, B., Goldberg, Y.P., Theilmann, J., Andrew, S.E., Zeisler, J., Adam, S., Greenberg, C., Ives, E.J., Clarke, L.A. et al. (1994) Somatic and gonadal mosaicism of the Huntington disease gene CAG repeat in brain and sperm. Nat Genet, 6, 409–414.

26. Tcw, J., Wang, M., Pimenova, A.A., Bowles, K.R., Hartley, B.J., Lacin, E., Machlovi, S.I., Abdelaal, R., Karch, C.M., Phatnani, H. et al. (2017) An Efficient Platform for Astrocyte Differentiation from Human Induced Pluripotent Stem Cells. Stem Cell Reports, 9, 600–614.

27. Sharp, P.M. and Li, W.H. (1987) The codon Adaptation Index--a measure of directional synonymous codon usage bias, and its potential applications. Nucleic Acids Res, 15, 1281–1295.

28. Lorenz, R., Bernhart, S.H., Honer Zu Siederdissen, C., Tafer, H., Flamm, C., Stadler, P.F. and Hofacker, I.L. (2011) ViennaRNA Package 2.0. Algorithms Mol Biol, 6, 26.

29. Richner, J.M., Himansu, S., Dowd, K.A., Butler, S.L., Salazar, V., Fox, J.M., Julander, J.G., Tang, W.W., Shresta, S., Pierson, T.C. et al. (2017) Modified mRNA Vaccines Protect against Zika Virus Infection. Cell, 169, 176.

30. Li, Y., Tenchov, R., Smoot, J., Liu, C., Watkins, S. and Zhou, Q. (2021) A Comprehensive Review of the Global Efforts on COVID-19 Vaccine Development. ACS Cent Sci, 7, 512–533.

31. Schoenmaker, L., Witzigmann, D., Kulkarni, J.A., Verbeke, R., Kersten, G., Jiskoot, W. and Crommelin, D.J.A. (2021) mRNA-lipid nanoparticle COVID-19 vaccines: Structure and stability. Int J Pharm, 601, 120586.

32. Lee, J.M., Zhang, J., Su, A.I., Walker, J.R., Wiltshire, T., Kang, K., Dragileva, E., Gillis, T., Lopez, E.T., Boily, M.J. et al. (2010) A novel approach to investigate tissue-specific trinucleotide repeat instability. BMC Syst Biol, 4, 29.

33. Keiser, M.S., Kordasiewicz, H.B. and McBride, J.L. (2016) Gene suppression strategies for dominantly inherited neurodegenerative diseases: lessons from Huntington’s disease and spinocerebellar ataxia. Hum Mol Genet, 25, R53–64.

34. Long, J.D., Lee, J.M., Aylward, E.H., Gillis, T., Mysore, J.S., Abu Elneel, K., Chao, M.J., Paulsen, J.S., MacDonald, M.E. and Gusella, J.F. (2018) Genetic Modification of Huntington Disease Acts Early in the Prediagnosis Phase. Am J Hum Genet, 103, 349–357.

35. Wild, E.J. and Tabrizi, S.J. (2017) Therapies targeting DNA and RNA in Huntington’s disease. Lancet Neurol, 16, 837–847.

36. Lee, J.M., Pinto, R.M., Gillis, T., St Claire, J.C. and Wheeler, V.C. (2011) Quantification of age-dependent somatic CAG repeat instability in Hdh CAG knock-in mice reveals different expansion dynamics in striatum and liver. PLoS One, 6, e23647.

37. Brandebura, A.N., Paumier, A., Onur, T.S. and Allen, N.J. (2023) Astrocyte contribution to dysfunction, risk and progression in neurodegenerative disorders. Nat Rev Neurosci, 24, 23–39.

38. Diaz-Castro, B., Gangwani, M.R., Yu, X., Coppola, G. and Khakh, B.S. (2019) Astrocyte molecular signatures in Huntington’s disease. Sci Transl Med, 11.

39. Rostalski, H., Leskela, S., Huber, N., Katisko, K., Cajanus, A., Solje, E., Marttinen, M., Natunen, T., Remes, A.M., Hiltunen, M. et al. (2019) Astrocytes and Microglia as Potential Contributors to the Pathogenesis of C9orf72 Repeat Expansion-Associated FTLD and ALS. Front Neurosci, 13, 486.

40. Skotte, N.H., Andersen, J.V., Santos, A., Aldana, B.I., Willert, C.W., Norremolle, A., Waagepetersen, H.S. and Nielsen, M.L. (2018) Integrative Characterization of the R6/2 Mouse Model of Huntington’s Disease Reveals Dysfunctional Astrocyte Metabolism. Cell Rep, 23, 2211–2224.

41. Shaltouki, A., Peng, J., Liu, Q., Rao, M.S. and Zeng, X. (2013) Efficient generation of astrocytes from human pluripotent stem cells in defined conditions. Stem Cells, 31, 941–952.

42. Chandrasekaran, A., Avci, H.X., Leist, M., Kobolak, J. and Dinnyes, A. (2016) Astrocyte Differentiation of Human Pluripotent Stem Cells: New Tools for Neurological Disorder Research. Front Cell Neurosci, 10, 215.

43. Zhou, Q., Viollet, C., Efthymiou, A., Khayrullina, G., Moritz, K.E., Wilkerson, M.D., Sukumar, G., Dalgard, C.L. and Doughty, M.L. (2019) Neuroinflammatory astrocytes generated from cord blood-derived human induced pluripotent stem cells. J Neuroinflammation, 16, 164.

44. Krencik, R., Weick, J.P., Liu, Y., Zhang, Z.J. and Zhang, S.C. (2011) Specification of transplantable astroglial subtypes from human pluripotent stem cells. Nat Biotechnol, 29, 528–534.

45. Krencik, R. and Zhang, S.C. (2011) Directed differentiation of functional astroglial subtypes from human pluripotent stem cells. Nat Protoc, 6, 1710–1717.

46. Colombo, E. and Farina, C. (2016) Astrocytes: Key Regulators of Neuroinflammation. Trends Immunol, 37, 608–620.

47. Diaz-Castro, B., Bernstein, A.M., Coppola, G., Sofroniew, M.V. and Khakh, B.S. (2021) Molecular and functional properties of cortical astrocytes during peripherally induced neuroinflammation. Cell Rep, 36, 109508.

48. Bates, G.P., Dorsey, R., Gusella, J.F., Hayden, M.R., Kay, C., Leavitt, B.R., Nance, M., Ross, C.A., Scahill, R.I., Wetzel, R. et al. (2015) Huntington disease. Nat Rev Dis Primers, 1, 15005.

49. Gonitel, R., Moffitt, H., Sathasivam, K., Woodman, B., Detloff, P.J., Faull, R.L. and Bates, G.P. (2008) DNA instability in postmitotic neurons. Proc Natl Acad Sci U S A, 105, 3467–3472.

50. Shelbourne, P.F., Keller-McGandy, C., Bi, W.L., Yoon, S.R., Dubeau, L., Veitch, N.J., Vonsattel, J.P., Wexler, N.S., Group, U.S.-V.C.R., Arnheim, N. et al. (2007) Triplet repeat mutation length gains correlate with cell-type specific vulnerability in Huntington disease brain. Hum Mol Genet, 16, 1133–1142.

51. Wheeler, V.C., Gutekunst, C.A., Vrbanac, V., Lebel, L.A., Schilling, G., Hersch, S., Friedlander, R.M., Gusella, J.F., Vonsattel, J.P., Borchelt, D.R. et al. (2002) Early phenotypes that presage late-onset neurodegenerative disease allow testing of modifiers in Hdh CAG knock-in mice. Hum Mol Genet, 11, 633–640.

52. Bradford, J., Shin, J.Y., Roberts, M., Wang, C.E., Sheng, G., Li, S. and Li, X.J. (2010) Mutant huntingtin in glial cells exacerbates neurological symptoms of Huntington disease mice. J Biol Chem, 285, 10653–10661.

53. Gillmore, J.D., Gane, E., Taubel, J., Kao, J., Fontana, M., Maitland, M.L., Seitzer, J., O’Connell, D., Walsh, K.R., Wood, K. et al. (2021) CRISPR-Cas9 In Vivo Gene Editing for Transthyretin Amyloidosis. N Engl J Med, 385, 493–502.

54. Lai, F., Hu, K., Wu, Y., Tang, J., Sang, Y., Cao, J. and Kang, T. (2012) Human KIAA1018/FAN1 nuclease is a new mitotic substrate of APC/C(Cdh1). Chin J Cancer, 31, 440–448.

55. McAllister, B., Donaldson, J., Binda, C.S., Powell, S., Chughtai, U., Edwards, G., Stone, J., Lobanov, S., Elliston, L., Schuhmacher, L.N. et al. (2022) Exome sequencing of individuals with Huntington’s disease implicates FAN1 nuclease activity in slowing CAG expansion and disease onset. Nat Neurosci, 25, 446–457.

56. Deshmukh, A.L., Caron, M.C., Mohiuddin, M., Lanni, S., Panigrahi, G.B., Khan, M., Engchuan, W., Shum, N., Faruqui, A., Wang, P. et al. (2021) FAN1 exo-not endo-nuclease pausing on disease-associated slipped-DNA repeats: A mechanism of repeat instability. Cell Rep, 37, 110078.

57. Alsuraih, M., LaViolette, B., Lin, G.-Y., Kovi, R., Daurio, N., Cheng, C., Ahn, Y., Jiang, Z., Ortiz, R., Li, S. et al. (2023) Targeting ABCB4 using mRNA-LNP for the treatment of rare liver diseases. bioRxiv, 2023.2004.2011.535868.

58. Olejniczak, M., Urbanek, M.O. and Krzyzosiak, W.J. (2015) The role of the immune system in triplet repeat expansion diseases. Mediators Inflamm, 2015, 873860.

59. Saba, J., Couselo, F.L., Bruno, J., Carniglia, L., Durand, D., Lasaga, M. and Caruso, C. (2022) Neuroinflammation in Huntington’s Disease: A Starring Role for Astrocyte and Microglia. Curr Neuropharmacol, 20, 1116–1143.

60. Creus-Muncunill, J. and Ehrlich, M.E. (2019) Cell-Autonomous and Non-cell-Autonomous Pathogenic Mechanisms in Huntington’s Disease: Insights from In Vitro and In Vivo Models. Neurotherapeutics, 16, 957–978.

61. Nakamori, M., Shimizu, H., Ogawa, K., Hasuike, Y., Nakajima, T., Sakurai, H., Araki, T., Okada, Y., Kakita, A. and Mochizuki, H. (2022) Cell type-specific abnormalities of central nervous system in myotonic dystrophy type 1. Brain Commun, 4, fcac154.

62. Smogorzewska, A., Desetty, R., Saito, T.T., Schlabach, M., Lach, F.P., Sowa, M.E., Clark, A.B., Kunkel, T.A., Harper, J.W., Colaiacovo, M.P. et al. (2010) A genetic screen identifies FAN1, a Fanconi anemia-associated nuclease necessary for DNA interstrand crosslink repair. Mol Cell, 39, 36–47.

63. Kratz, K., Schöpf, B., Kaden, S., Sendoel, A., Eberhard, R., Lademann, C., Cannavó, E., Sartori, A.A., Hengartner, M.O. and Jiricny, J. (2010) Deficiency of FANCD2-Associated Nuclease KIAA1018/FAN1 Sensitizes Cells to Interstrand Crosslinking Agents. Cell, 142, 77–88.

64. Liu, T., Ghosal, G., Yuan, J., Chen, J. and Huang, J. (2010) FAN1 acts with FANCI-FANCD2 to promote DNA interstrand cross-link repair. Science, 329, 693–696.

65. Cannavo, E., Gerrits, B., Marra, G., Schlapbach, R. and Jiricny, J. (2007) Characterization of the interactome of the human MutL homologues MLH1, PMS1, and PMS2. J Biol Chem, 282, 2976–2986.

