## Supplemental Information for "Increased FAN1 expression by mRNA-LNP attenuates CAG repeat expansion in Huntington patients’ iPSC-derived astrocytes"

**Supplemental Figure S1:**

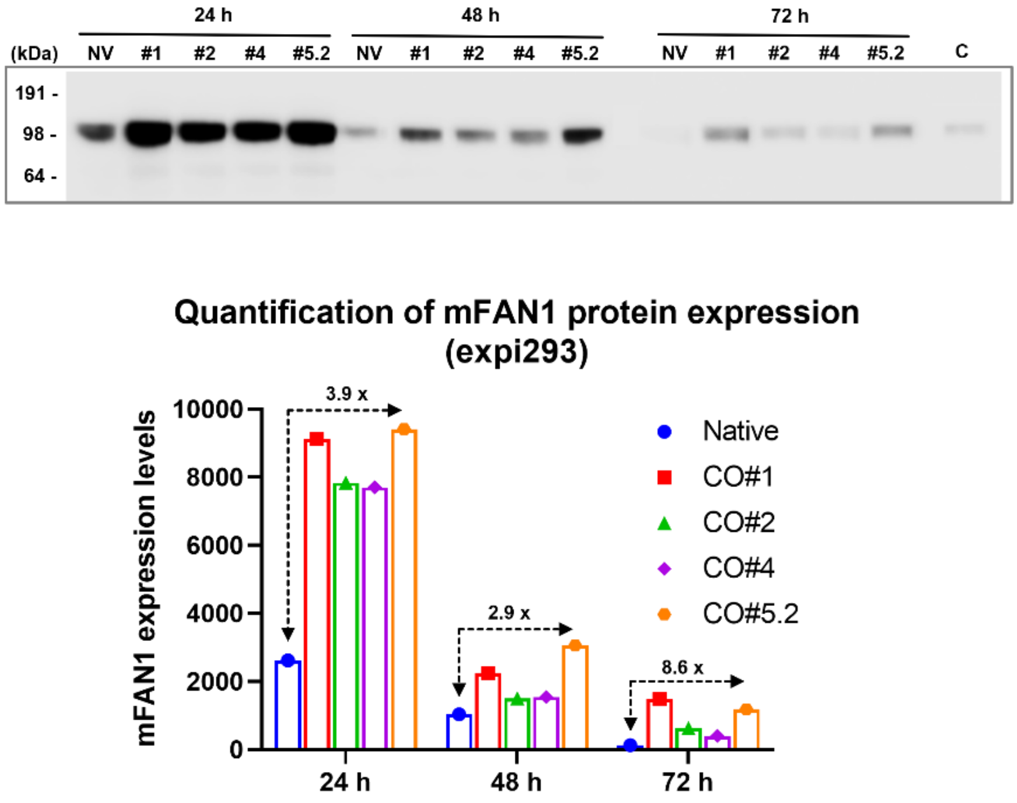


**Figure S1. Characterization of native and codon-optimized *mFan1* mRNAs in expi293 cell line.** Western blot monitoring of FAN1 protein expression in expi293 cell line transfected with *mFan1* mRNA of native (NV) coding sequence or different codon-optimized coding sequences *(#1, #2, #4, and #5.2)* encoding murine FAN1 proteins (24, 48 or 72 hrs after transfection). Positive control (C): recombinant mouse FAN1 protein. Bottom panel: Quantification of FAN1 protein expression in expi293 cell line after transfection with codon-optimized *mFan1* mRNAs shown in top panel.

### Supplemental Table S1: Weights of synonymous replacement codons

| Amino Acid | Codon | Relative Weight |
|---|---|---|
| Ala | GCA | 21 |
| Ala | GCC | 40 |
| Ala | GCG | 7 |
| Ala | GCT | 32 |
| Arg | AGA | 18 |
| Arg | AGG | 18 |
| Arg | CGA | 11 |
| Arg | CGC | 24 |
| Arg | CGG | 17 |
| Arg | CGT | 11 |
| Asn | AAC | 60 |
| Asn | AAT | 40 |
| Asp | GAC | 53 |
| Asp | GAT | 46 |
| Cys | TGC | 62 |
| Cys | TGT | 38 |
| Gln | CAA | 23 |
| Gln | CAG | 77 |
| Glu | GAA | 37 |
| Glu | GAG | 63 |
| Gly | GGA | 27 |
| Gly | GGC | 34 |
| Gly | GGG | 19 |
| Gly | GGT | 20 |
| His | CAC | 62 |
| His | CAT | 37 |
| Ile | ATA | 12 |
| Ile | ATC | 52 |
| Ile | ATT | 36 |
| Leu | CTA | 9 |
| Leu | CTC | 20 |
| Leu | CTG | 40 |
| Leu | CTT | 14 |
| Leu | TTA | 5 |
| Leu | TTG | 12 |
| Lys | AAA | 35 |
| Lys | AAG | 65 |
| Met | ATG | 100 |
| Phe | TTC | 56 |
| Phe | TTT | 44 |
| Pro | CCA | 28 |

| | | |
|---|---|---|
| Pro | CCC | 34 |
| Pro | CCG | 5 |
| Pro | CCT | 33 |
| Thr | ACA | 26 |
| Thr | ACC | 41 |
| Thr | ACG | 7 |
| Thr | ACT | 26 |
| Trp | TGG | 100 |
| Tyr | TAC | 59 |
| Tyr | TAT | 41 |
| Val | GTA | 10 |
| Val | GTC | 24 |
| Val | GTG | 47 |
| Val | GTT | 18 |

**Supplemental Table S2: Optimization schedule for Method 1**

| | | | | | | | Optimization Metric Weights | | | |
|---|---|---|---|---|---|---|---|---|---|---|
| Round Type | CPUs | Generations | Child sequences per generation | Initial mutation rate | Mutation rate decay per generation | Sequences passed to next round | CAI | Rolling CAI | MFE Unpaired Fraction | Long Run Fraction |
| SD | 1 | NA | NA | NA | NA | 1 | 1 | 0 | 0 | 0 |
| MC/GA | 50 | 50 | 20 | 0.3 | 0.002 | 20 | -1 | 0 | 0 | 0 |
| MC/GA | 40 | 20 | 100 | 0.1 | 0.002 | 20 | -1 | 0 | 0 | 0 |
| SD | 20 | NA | NA | NA | NA | 20 | -1 | 0 | 0 | 0 |
| MG/GA | 40 | 10 | 10 | 0.05 | 0.005 | 10 | -1 | -0.1 | 1 | -0.1 |

**Supplemental Table S3: Optimization schedule for Method 2**

| | | | | | | | Optimization Metric Weights | | | |
|---|---|---|---|---|---|---|---|---|---|---|
| Round Type | CPUs | Generations | Child sequences per generation | Initial mutation rate | Mutation rate decay per generation | Sequences passed to next round | CAI | Rolling CAI | MFE Unpaired Fraction | Long Run Fraction |
| SD | 1 | NA | NA | NA | NA | 1 | 1 | 0 | 0 | 0 |
| MC/GA | 50 | 50 | 20 | 0.3 | 0.002 | 40 | -1 | 0 | 0 | 0 |
| MC/GA | 40 | 20 | 100 | 0.1 | 0.002 | 20 | -1 | 0 | 0 | 0 |
| SD | 20 | NA | NA | NA | NA | 1 | -1 | 0 | 0 | 0 |
